# The Rotating Snake Illusion is a straightforward consequence of non-linearity in arrays of standard motion detectors

**DOI:** 10.1101/2020.04.30.070623

**Authors:** Michael Bach, Lea Atala-Gérard

## Abstract

The Rotating Snakes illusion is a motion illusion based on repeating, asymmetric luminance patterns. Recently, we found certain grey-value conditions where a weak, illusory motion occurs in the opposite direction. Of the four models for explaining the illusion, one (Backus and Oruç, 2005) also explains the unexpected perceived opposite direction. We here present a simple new model, without free parameters, based on an array of standard correlation-type motion detectors with a subsequent non-linearity (e.g., saturation) before summing the detector outputs. The model predicts (1) the pattern-appearance motion illusion for steady fixation, (2) an illusion under the real-world situation of saccades across or near the pattern (pattern shift), (3) a relative maximum of illusory motion for the same grey values where it is found psychophysically, and (4) the inverse illusion for certain luminance values. We submit that the model’s sparseness of assumptions justifies adding a fifth model to explain this illusion.

## Introduction

Certain spatial patterns can evoke illusory movement, especially under dynamic viewing. An early example was the Peripheral Drift Illusion (Fraser & Wilcox, 1979). A. Kitaoka optimized its patterns and colors, leading to the strong, beautiful and widely known “Rotating Snakes Illusion” (Kitaoka (2003; Kitaoka & Ashida, 2003, Fig. 5). While usually rendered in color, it is nearly just as strong in grey (Conway et al., 2005). The Rotating Snakes Illusion with its many variations continues to fascinate beyond the vision research community. It basically needs four luminance levels in an asymmetric spatial arrangement, e.g. black/light-grey/dark-grey/white (Murakami et al., 2006). When we assessed the optimal luminance conditions for the intermediate grey levels, we found an as yet unknown “parameter island of weak opposite rotation” when mapped into the plane of light-grey vs. dark grey values for the middle two patches (Atala-Gérard & Bach, 2017).

To date, there are four models explaining the illusory motion in the Rotating Snakes Illusion (Backus & Oruç, 2005; Conway et al., 2005; Fermüller et al., 2010; Murakami et al., 2006). As we have discussed previously, only the Backus and Oruç (2005) model is able to predict the “island of opposite rotation” (Atala-Gérard & Bach, 2017). However, for this to work it requires a specific contrast transfer function (Atala-Gérard, 2018) which differs from the one used by Backus and Oruç (2005). Furthermore, in the natural viewing situation, the Snake pattern does not suddenly appear from neutral background, as assumed in this model, and for seeing the illusion, saccades are necessary (small or large (Otero-Millan et al., 2012)).

We here present a fifth, simpler and parameter-free model, based on nothing but an array of standard Reichard correlation detectors [which are equivalent to the motion energy model (Adelson & Bergen, 1985), but see (Borst, 2007)] with a subsequent non-linear (e.g. saturating) transfer function before summing their outputs. This, together with saccades while viewing, or the appearance of the pattern out of a grey background, predicts the standard Rotating Snakes Illusion, including a parameter region leading to the opposite direction of rotation.

These findings suggest that the Rotating Snakes Illusion can be regarded as a necessary side effect when arrays of motion detectors are combined in a non-linear fashion.

### A new, simple computational model

We will present the model in two steps, beginning with a simpler situation, namely that of pattern-appearance of the stimulus from a grey background. We will then move on to treat natural viewing conditions, namely that the observer performs saccades across the stimulus picture. The model only assumes the presence of arrays (Snippe & Koenderink, 1994) of standard Reichardt-Hassenstein correlation detectors (Borst & Egelhaaf, 1989; Hassenstein & Reichardt, 1956, Fig. 4e; Reichardt, 1986). As these are mathematically equivalent with the motion energy model (Adelson & Bergen, 1985), all our findings will hold for the energy model as well. In the model presented here, it proved necessary to add a sign-conserving non-linearity (of nearly any shape, see below) at the output of individual motion detectors, before averaging across the detector array. Saturating non-linearities are frequently observed in neural systems (Peirce, 2007) and, specifically, in the motion system (Derrington & Goddard, 1989) and were suggested by (Adelson & Bergen, 1985): “A compressive nonlinearity (such as a square root) may follow…”. We tested various sigmoid functions that all share the property of being rotationally symmetric around zero, including the arc tangent, hyperbolic tangent, logistic function.

**Fig. 1.**
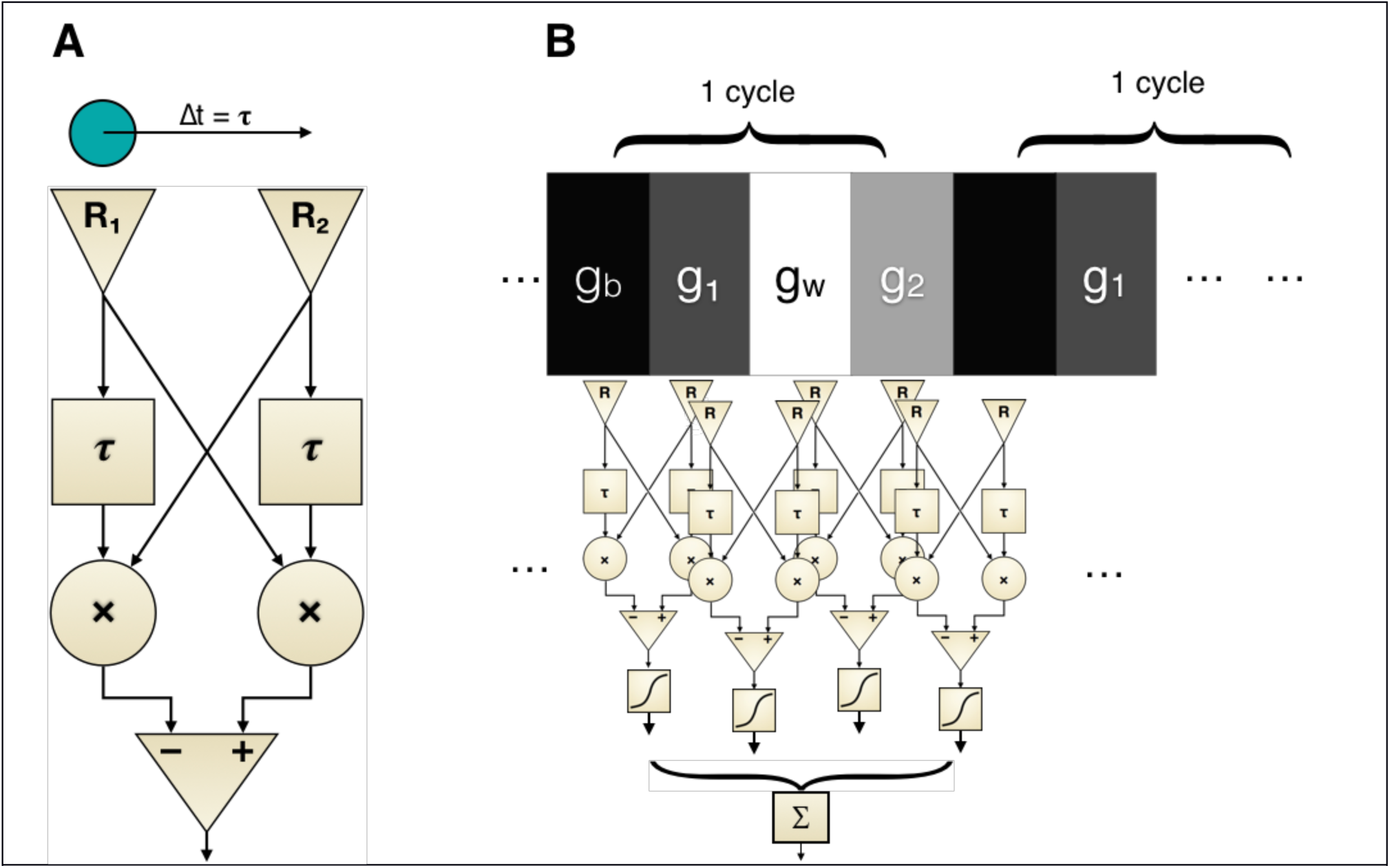
**A**. Standard correlation-type motion detector (Borst & Egelhaaf, 1989; Reichardt, 1987). The stimulus (circle at top) moves from receptor R_1_ to R_2_ in time Δt, which is identical to the delay stages’ time constant τ for a maximal response of the detector. After multiplication of the delayed with the non-delayed receptor output at “×”, the signals from the two paths are subtracted, resulting in a bipolar output on left/rightward motion, and a suppressed flicker response. B. Two sample cycles of a Snakes sequence (Murakami et al., 2006) supplied as an input to an array of motion detectors, each detector spatially tuned to the distance of a Snakes-sequence cell. The output of each individual motion detector passes a non-linear transfer function, before being summed across the array of motion detectors, yielding a net motion output.

**Fig. 2.**
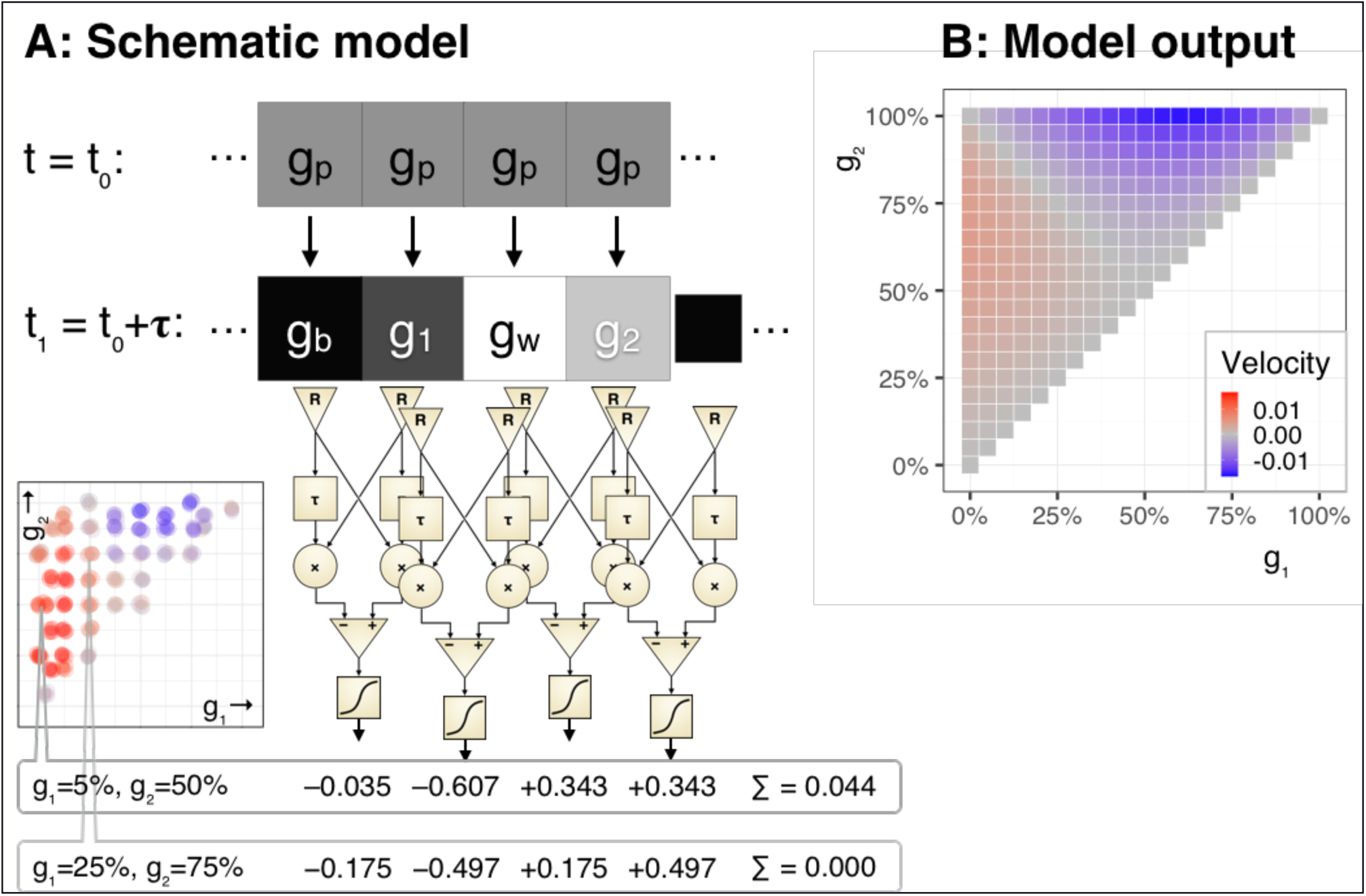
Model, first approximation. **A**. On top, a sequence of grey squares turns into the Snake sequence below. That sequence is the input to an array of motion detectors (cf. Fig. 1), whose output is calculated for two example combinations of (g_1_, g_2_). The positions of these (g_1_, g_2_) are indicated on a psychophysical data map (bottom left) which we reported previously (Atala-Gérard & Bach, 2017), showing two parameter regions which result in opposite illusory motion direction. The first (g_1_, g_2_) pair is from the region of strongest observed illusion, and indeed the average of the four outputs yields a non-negative value. The second (g_1_, g_2_) pair (25%, 75%) is from the “valley” between the two regions (the diagonal), and for symmetry reasons yields a zero net motion output. **B**. Net motion output example for all (g_1_, g_2_) pairs. It bears a qualitative similarity to the psychophysical findings (shown in figure part A, bottom left): There is a motion maximum near the expected region (≈5%, 50%) and, indeed, the model predicts an inverse direction near the region where this was experimentally observed (top right).

**Fig. 3.**
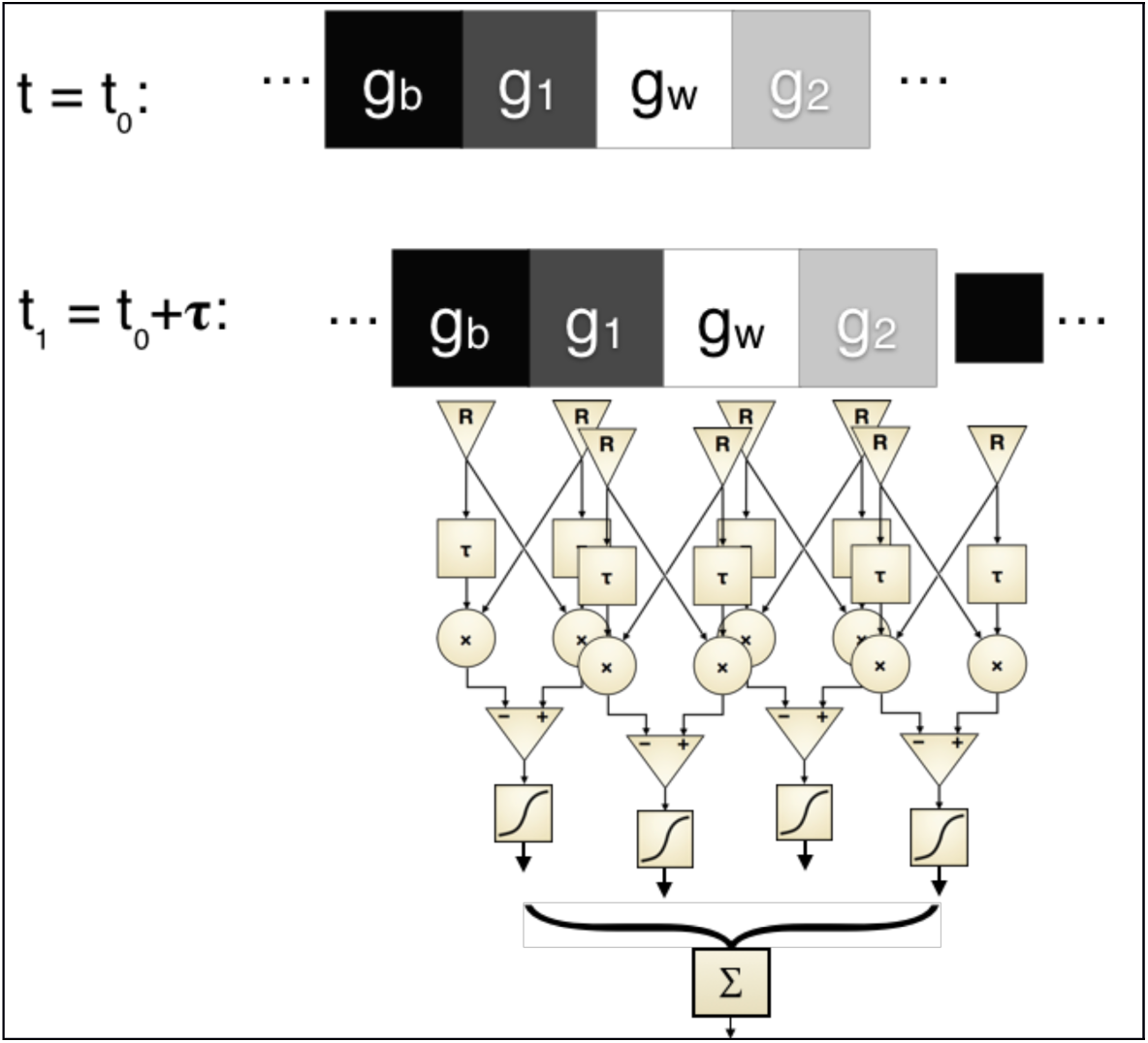
The simplification of the previous figure (pattern appearance: Snakes appearing from grey) is now made more realistic: Snakes appearing from all possible horizontal shifts of a previous Snake pattern. The shift shown for (half a cell width) is but one example of the possible shifts. The motion detectors are arranged exactly as in the first approximation.

**Fig. 4.**
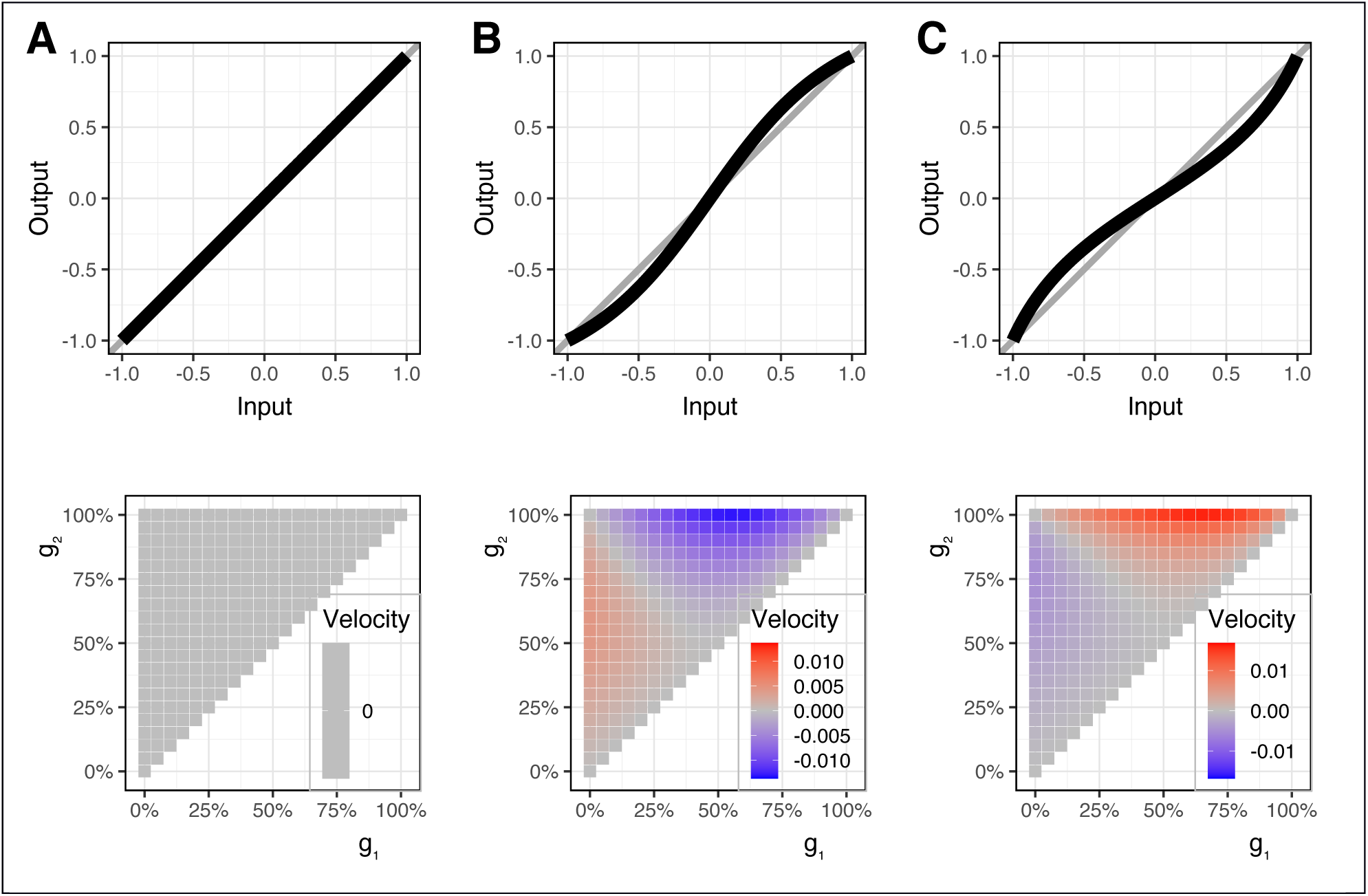
Top: Three different transfer functions (A, linear; B, tangent; C, arc tangent). Bottom: the net output of the random-position appearance model for all possible (g_1_, g_2_) combinations. **A**. With a linear transfer function, no net motion signal appears. **B**. We tested a range of saturating non-linearities and in all cases there appears a net motion signal for any off-diagonal (i.e., asymmetrical) (g_1_, g_2_) combination, and also a polarity inversion when crossing the diagonal – as also found psychophysically. **C**. Accelerating non-linearities also lead to a net motion output, with an inverted sign (direction) of apparent motion compared to saturating non-linearities.

#### First approximation: pattern appearance

This first approximation to a working model is based on the observation that the appearance of a Snake Pattern from a grey background (pattern appearance) evokes a strong apparent motion even with steady fixation. This is demonstrated on the website https://michaelbach.de/ot/mot-snakesLum/ (Bach, 2020a) if “Modulate contrast” is selected there. To make the geometry more tractable, we uncoiled the original Rotating Snake Illusion with its several “Snake wheels”, and investigated a pattern consisting of repeated “Snake cycles”, each cycle containing four grey values (Fig. 1B).

Basic motion detectors (Fig. 1A, based on Reichardt (1987, his Fig. 4c) include a delay τ, which we account for by simply comparing the time points before and after appearance (t_0_, t_1_, respectively) thus leaving the exact delay time undetermined (which may be 50–100 ms). The spatial span of each detector R equals the width of one stripe of the Snake pattern, and there is no space between adjacent detectors. The sensitivity of the detectors is spatially constant and normalized to 1 so that the output of the detectors is simply *∫ L* (*x*)*dx*, with L(x) being the luminance and the limits of the integral the spatial span of the respective detector. We use the sign convention that a dark structure on light background yields a positive output when moving to the right. Given

*g*_1_: grey value at t_1_ at the right input,

*g*_1*p*_: previous (t_0_) grey value at the right input,

*g*_2_: grey value at t_1_ at the left input,

*g*_2*p*_: previous (t_0_) grey value at the left input,

and assuming some function *f* (which could be the identity),

the output *d* of the Reichard detector can be calculated by

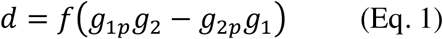

##### Main assumption

The sum (or the average, these differ here only by a scaling factor) of an array of such motion detectors, stimulated by the appearance of a Snake cycle, subserves the apparent motion (Fig. 1B). Thus, given

*g*_*p*_: the previous (at t_0_) background grey value at all inputs,

*g*_*b*_ and *g*_*w*_ for the black and white value, respectively,

the sum *d_∑_* of four detectors, stimulated by a full Snake cycle, will be given by

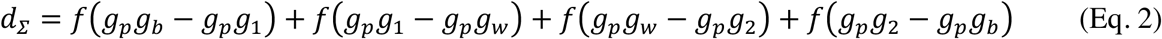

Note that *d*_*∑*_ will collapse to zero if ***f*** is the identity function. For our purposes we will only require ***f*** to be a mapping from [-1, 1] to [-1, 1], point-symmetrical around zero, with *f*(0) = 0, *f*(1) = 1 and *f*(−1) = −1. In this generality, we could not solve the problem analytically, so implemented it as a computational model in the R language (R Core Team, 2020), a free open-source programming and statistical environment, and graphs were produced using the package ggplot2. [Full source code in the repository (Bach, 2020b)].

Fig. 2**A** shows the motion detector array along a Snake cycle with two examples of grey-value pairs (bottom left), and corresponding summed outputs ∑. To give but two examples, for the combination (g_1_, g_2_) = (0.25, 0.75), the net output was zero; for the combination (g_1_, g_2_) = (0.05, 0.5), a non-zero output resulted. Non-zero net outputs only occurred with the insertion of the aforementioned non-linearity.

Fig. 2B shows the net motion for the full parameter space of the possible grey values (g_1_, g_2_). We observed a maximum velocity around the region where that was expected (g_1_≈5%, g_2_≈50%) and, indeed, found opposite motion direction where the psychophysical data predict opposite direction of illusory motion.

The various shapes of the non-linearity affected the magnitude of the net motion output, but the distribution in (g_1_, g_2_) space remained the same (for all the tested non-linearities). Non-linearities within the receptors themselves were tested as well, but had no qualitative effect in this model and were thus omitted from further analysis, although physiologically they are likely to occur.

#### Closer approximation, full model: “Pattern shift”, appearance at random positions along the Snake cycle

We model the effects of saccades across the image by stimulating the motion detector array with Snake cycles changing their positions randomly (Fig. 3). Due to Kitaoka’s (2003; Kitaoka & Ashida, 2003, Fig. 5) arrangement in Snake wheels, saccades can affect the position of a Snake cycle as a random shift (Fig. 3) or at any other angle. Assuming that the latter average out, we will only compute the effects of lateral shifts here.

In Fig. 3, we sketch the model structure: As an initial condition, a Snakes sequence could have any possible shift relative to the final position which is equal to the position of the detectors R_i_. Since the problem is circular, we averaged over 40 small shifts of the Snakes sequence until it was identical again. The results are shown in Fig. 4, using three different non-linearities. Unsurprisingly, no motion illusion appears for a linear transfer function (A). For any (of the tested) saturating transfer functions a non-zero (illusory) motion, including the “opposite island”, occurs with the pattern shift model just as with the pattern appearance model. Fig. 4C adds an accelerating non-linearity: illusory motion appears again, but with opposite sign; the same sign reversal occurs also in the pattern-appearance model (not depicted).

For *all* tested non-linearities, we found areas in the (g_1_, g_2_) plane which give non-zero results (and thus illusory motion), but the relative areas of positive and negative motion varied slightly.

## Discussion

We present a computational model which demonstrates that arrays of standard motion detectors (of the correlation- or motion-energy type), followed by a non-linear transfer function, exhibit the motion illusion known as the Rotating Snakes illusion. The non-linearity can have nearly any shape, as long as it is monotonous. Some properties of psychophysical findings are predicted by the model: (1) prediction of the pattern-appearance motion illusion for steady fixation, (2) an illusion under the natural viewing situation of performing saccades across the pattern (pattern shift), (3) the presence of a relative maximum of illusory motion right at the location in the (g_1_, g_2_) parameter space where it is found psychophysically, and (4) the (recently discovered) inverse illusion in a certain parameter region. There are a number of shortcomings and assumptions associated with our simple model which will be discussed next.

While several aspects of the illusion are captured by the model, there is no *quantitative* prediction, and it predicts equal strength for the “island of opposite rotation” which perceptually is markedly weaker.

Initially we had considered that the real-world saccadic condition can also be traced to pattern-appearance, invoking saccadic suppression to transform pattern-shift into pattern appearance. This may well be the case, but it seems that saccadic suppression is not needed to explain the illusion.

The most famous version of the “Rotating Snakes” (Kitaoka, 2003) is in color, which we have simplified here to a luminance-only version. This is, of course, computationally much more tractable and the simplification seems justified, as Conway et al. (2005) found similar illusion strengths for the luminance and color variants.

While the present model needs no free parameters to be fitted, there are a number of inherent simplifying assumptions: A major one is that the spatial tuning of the detector inputs being matched to the Snake pattern sequence (Fig. 1B). This may indeed be related to the finding that the illusion is typically strongest when performing saccades in the neighborhood of the picture. Consequently, somewhat larger receptive fields (Strasburger et al., 2011) may become involved, that are spatially better tuned to the pattern. Furthermore, optimal matches between the stimulus and the motion detector arrangement will be only fleeting, which matches the perception of this illusion. The model rests on summation of motion receptors, thus loosing spatial information in the model. But some aggregation of motion detectors is required anyway to account e.g. for higher-order motion (Lu & Sperling, 1996) or independence of form (Glünder, 1990).

A very specific shortcoming of the model is the missing prediction of the factual illusion weakness in what we call the “island of opposite rotation” as opposed to the standard grey-value region (it rather predicts the same strength). A further assumption is connected to our simplified saccade model. In the model, we consider the saccade’s motion trajectory as an instantaneous step function. However, in reality, saccades will occur more or less randomly when viewing the Snake patterns. Thus, our “pattern shift” situation is only one of many possibilities that can occur. Our assumption that all other angles and positions will “average out” may well deserve more detailed scrutiny.

While the above is a long, and possibly incomplete, list of assumptions, they all appear physiologically plausible. We were surprised that this very simple model predicts more properties of the Rotating Snakes illusion than any of the previous models, and that it yielded similar results for any of the tested, monotonous non-linearities.

## Conclusion

We demonstrate that an array of standard motion detectors with a non-linear transfer function for each detector before summing the individual receptor outputs gives rise to a motion signal, which qualitatively shows all the known properties of the “Rotating Snakes” illusion. We submit that more complicated models are not required to explain this illusion, since it appears to be a straightforward consequence of the non-linearity (which is widely found in the nervous system) when confronted with the repeated, spatially asymmetric grey-value sequence of the Rotating Snakes illusion. Taken together, this underlines the notion that understanding the mechanisms of illusion can be an automatic byproduct of understanding mechanisms of general visual perception.

## Acknowledgements

We thank Hans Strasburger for very careful correction of an earlier version of the manuscript.

The article processing charge was funded by the Baden-Wuerttemberg Ministry of Science, Research and Art and the University of Freiburg in the funding program “Open Access Publishing”.

## References

Adelson, E. H., & Bergen, J. R. (1985). Spatiotemporal energy models for the perception of motion. JOSA A, 2(2), 284–299. https://doi.org/10.1364/JOSAA.2.000284

Atala-Gérard, L. (2018). Quantitative Analyse der Bewegungstäuschung “Rotating Snakes.” Thesis. https://doi.org/10.6094/UNIFR/16964

Atala-Gérard, L., & Bach, M. (2017). Rotating Snakes Illusion – Quantitative analysis reveals a region in luminance space with opposite illusory rotation. I-Perception, 8(1), 2041669517691779. https://doi.org/10.1177/2041669517691779

Bach, M. (2020a). “Rotating Snakes” with luminance control. https://michaelbach.de/ot/mot-snakesLum/

Bach, M. (2020b). Rotating Snakes Illusion – Simple computational model. https://doi.org/10.6084/m9.figshare.12212666.v1

Backus, B. T., & Oruç, İ. (2005). Illusory motion from change over time in the response to contrast and luminance. Journal of Vision, 5(11), 10–10. https://doi.org/10.1167/5.11.10

Borst, A. (2007). Correlation versus gradient type motion detectors: The pros and cons. Philosophical Transactions of the Royal Society B: Biological Sciences, 362(1479), 369–374. https://doi.org/10.1098/rstb.2006.1964

Borst, A., & Egelhaaf, M. (1989). Principles of visual motion detection. Trends in Neurosciences, 12(8), 297–306. https://doi.org/10.1016/0166-2236(89)90010-6

Conway, B. R., Kitaoka, A., Yazdanbakhsh, A., Pack, C. C., & Livingstone, M. S. (2005). Neural basis for a powerful static motion illusion. Journal of Neuroscience, 25(23), 5651–5656. https://doi.org/10.1523/JNEUROSCI.1084-05.2005

Derrington, A. M., & Goddard, P. A. (1989). Failure of motion discrimination at high contrasts: Evidence for saturation. Vision Research, 29(12), 1767–1776. https://doi.org/10.1016/0042-6989(89)90159-4

Fermüller, C., Ji, H., & Kitaoka, A. (2010). Illusory motion due to causal time filtering. Vision Research, 50(3), 315–329. https://doi.org/10.1016/j.visres.2009.11.021

Fraser, A., & Wilcox, K. J. (1979). Perception of illusory movement. Nature, 281(5732), 565–566. https://doi.org/10.1038/281565a0

Glünder, H. (1990). Correlative velocity estimation: Visual motion analysis, independent of object form, in arrays of velocity-tuned bilocal detectors. JOSA A, 7(2), 255–263. https://doi.org/10.1364/JOSAA.7.000255

Hassenstein, B., & Reichardt, W. (1956). Systemtheoretische Analyse der Zeit-, Reihenfolgen- und Vorzeichenauswertung bei der Bewegungsperzeption des Rüsselkäfers Chlorophanus. Zeitschrift Für Naturforschung B, 11(9–10), 513–524. https://doi.org/10.1515/znb-1956-9-1004

Kitaoka, A. (2003). Rotating Snakes. http://www.psy.ritsumei.ac.jp/akitaoka/rotsnakee.html

Kitaoka, A., & Ashida, H. (2003). Phenomenal characteristics of the peripheral drift illusion. VISION: The Journal of the Vision Society of Japan, 15, 261–262. https://doi.org/10.24636/vision.15.4_261

Lu, Z.-L., & Sperling, G. (1996). Three Systems for Visual Motion Perception. Current Directions in Psychological Science, 5(2), 44–53. JSTOR. https://www.jstor.org/stable/20182388

Murakami, I., Kitaoka, A., & Ashida, H. (2006). A positive correlation between fixation instability and the strength of illusory motion in a static display. Vision Research, 46(15), 2421–2431. https://doi.org/10.1016/j.visres.2006.01.030

Otero-Millan, J., Macknik, S. L., & Martinez-Conde, S. (2012). Microsaccades and Blinks Trigger Illusory Rotation in the “Rotating Snakes” Illusion. The Journal of Neuroscience, 32(17), 6043–6051. https://doi.org/10.1523/JNEUROSCI.5823-11.2012

Peirce, J. W. (2007). The potential importance of saturating and supersaturating contrast response functions in visual cortex. Journal of Vision, 7(6), 13–13. https://doi.org/10.1167/7.6.13

R Core Team. (2020). R: A Language and Environment for Statistical Computing [R]. R Foundation for Statistical Computing. http://www.R-project.org

Reichardt, W. (1986). Processing of optical information by the visual system of the fly. Vision Research, 26(1), 113–126. https://doi.org/10.1016/0042-6989(86)90075-1

Reichardt, W. (1987). Evaluation of optical motion information by movement detectors. Journal of Comparative Physiology A, 161(4), 533–547. https://doi.org/10.1007/BF00603660

Snippe, H. P., & Koenderink, J. J. (1994). Extraction of optical velocity by use of multi-input Reichardt detectors. JOSA A, 11(4), 1222–1236. https://doi.org/10.1364/JOSAA.11.001222

Strasburger, H., Rentschler, I., & Jüttner, M. (2011). Peripheral vision and pattern recognition: A review. Journal of Vision, 11(5), 13–13. https://doi.org/10.1167/11.5.13

